# Androgen receptor plays critical role in regulating cervical cancer cell migration

**DOI:** 10.1101/2024.07.13.603408

**Authors:** Sarpita Bose, Subhrangshu Das, Sebabrata Maity, Oishee Chakrabarti, Saikat Chakrabarti

## Abstract

Cervical cancer (CC) is the second most common cancer among women in India and the fourth worldwide. While major genes and pathways have been studied, further research is needed to identify candidates for targeted therapy in metastatic disease. This study used a network biology approach to identify key genes in disease progression. Stage-specific cervical cancer protein-protein interaction networks (PPIN) were constructed by overlaying stage-specific, patient-derived transcriptomics data onto a human protein-protein interaction network (HPPIN). Graph-theory-based network analysis identified important interacting proteins (IIPs) with maximum connectivity, high centrality scores, and significant global and local network perturbation scores. Among the identified IIPs, the Androgen receptor (AR) emerged as one of the crucial yet understudied regulator in cervical cancer. Patient samples and in vitro experiments showed significant downregulation in cervical cancer. Ligand-dependent overexpression of AR reduced cancer cell migration while failed to induce apoptosis in CC cell lines. Downregulation of mesenchymal markers and restoration of epithelial markers suggested AR’s potential in reversing invasive properties of cervical cancer cells. AR overexpression upregulated its downstream target PTEN and restored GSK3β activity by interfering with AKT phosphorylation, probably leading to degradation of mesenchymal markers. Further studies showed AR reduced cell motility by hindering focal adhesion formation and Actin filament assembly. An increased G-Actin ratio suggested AR disrupted cytoskeletal dynamics through the RhoA/ROCK1/LIMK1/CFL1 pathway, impeding cervical cancer cell spread.

## Introduction

Cervical cancer (CC) is a significant health issue globally, ranking fourth in terms of both incidence and mortality worldwide. However, its impact in India is particularly pronounced, where it is the second most prevalent malignancy affecting women. Several factors contribute to the high burden of cervical cancer in India, including lower socioeconomic status, limited access to healthcare resources, and a high prevalence of human papillomavirus (HPV) infection [1]. In India, cervical cancer contributes to approximately 6–29% of all cancers in women [2]. Persistent infection with high-risk human papillomavirus (HPV) is a critical factor in the genesis and development of cervical cancer. While HPV vaccines could be a preventive measure, in its metastatic state, cervical cancer is indeed considered incurable with current treatment options; this emphasizes the need for targeted therapy [3]. It is a complex disease involving hundreds of pathways for disease progression. It is obvious that studying all or multiple of these pathways simultaneously is experimentally quite challenging; hence aptly underlying the necessity for application of computational biology in cervical cancer research as a fast, reliable and holistic platform.

This study utilizes a multi-pronged approach to first employ systems biology in utilizing protein-protein interaction data to comprehend and identify functionally significant proteins in human cancer. Protein-protein interaction networks are constructed and analyzed to investigate disease-specific scenarios [4–6]. These biological networks are found to follow the principles of graph and information theory [7]. Certain proteins have been designated as important interacting proteins (IIPs) based on graph theory principles. Nodes with maximum connectivity, higher centrality scores, and elevated global and local perturbation scores are regarded as significant within a biological network. A node meeting more than one of these criteria is classified as an IIP [8]. The application of this network biological approach may reveal functionally significant targets that play critical roles in the occurrence and progression of cervical cancer. A comparative study of transcriptomics data obtained from normal and cancerous tissue aids in constructing the supra-interaction network. Mathematical model-based network analysis facilitates the identification of key supra-regulatory network proteins that likely play crucial roles in cancer development and progression. Finally, *in silico* perturbation analysis and experimental validation may lead to the identification of novel connections and involved genes/proteins, which can be further considered as diagnostic and prognostic markers. This comprehensive approach integrates high-confidence protein-protein interactions with transcriptomics data to provide a robust foundation for exploring molecular interactions and regulatory processes within the context of cervical cancer.

Here, in this study Androgen receptor (AR) has emerged as a candidate target in cervical cancer through network biology analyses. AR is part of the steroid receptor superfamily, which governs gene expression in a ligand-dependent manner. Typically found in the cytoplasm when devoid of androgens, AR relocates to the nucleus upon binding to its ligand 5α-dihydrotestosterone (DHT), to initiate gene expression. As a transcription factor, AR is involved in regulating the expression of genes that control a wide range of biological processes including cell proliferation, migration, and apoptosis. AR regulates the transcriptional activity and expression of downstream target genes by interacting with the coactivators or corepressors, as well as upstream or downstream regulators through classical or non-classical AR signaling pathways [9]. Although AR is a well-established marker for prostate cancer, its role in gynecological cancers, specifically cervical cancer, remains largely unknown [10]. There are only a couple of studies that have shown downregulation of AR in cervical cancer, though without providing any mechanistic details [11, 12].

Here, through computational systems biology approach we first postulate a probable important role of AR in development as well as progression of cervical cancer. Expressions of AR both at mRNA and protein levels were found to be significantly downregulated in samples from patients, as well as in CC-derived cell lines, like HeLa and SiHa. Subsequently, to gain mechanistic insight into the role played by AR in the pathogenesis of cervical cancer progression, we undertook various *ex vivo* studies. Our results show that exogenous expression of AR followed by ligand (DHT) activation are essential for inhibition of migration and reduction of invasive potential by suppressing the mesenchymal markers essential for cervical cell migration. AR transcriptionally activated PTEN thereby restoring GSK3β activity. GSK3β facilitates proteasomal degradation of Snail, a transcriptional repressor of the epithelial marker E-cadherin. Further we observed increased E-cadherin expression indicative of restored epithelial properties. We also report that exogenous expression and ligand activation of AR make CC cells less motile with the presence of smaller focal adhesions. AR also destabilizes the Actin network via RhoA/ROCK1/LIMK1/CFL1 pathway, uncovering a novel mechanism of AR in cervical cancer. To the best of our knowledge, this is the first report highlighting the plausible mechanisms by which higher abundance and activation of AR could affect cervical cancer progression. Our study leads to avenues for future research in exploring DHT or synthetic androgens as potential therapeutic agents in treatment options for cervical cancer.

## Materials and Methods

### Data Collection

Human protein-protein interaction (HPPIN) data was extracted from the STRING interactome database, using protein-protein interactors established by high-throughput analyses with a high confidence score (≥0.7) to generate a protein-protein interaction network (PPIN) [13, 14]. Further, transcriptomics data were sourced from the NCBI GEO database [15]. A cervical cancer-specific transcriptomics dataset (GSE9750) was selected, comprising a total number of 66 samples, among which 33 are primary tumors, 9 are cell lines, and the remaining 24 are normal cervical epithelium samples.

### Construction of stage-specific PPIN

The raw data from 33 tumor samples (5 Stage I, 11 Stage II, 10 Stage III and1 Stage IV) and 2 cell lines (HeLa & SiHa) were normalized against corresponding normal samples utilizing the GEO2R analyzer. Due to insufficient samples in Stage IV, Stage III and IV data were merged. The normalized data underwent further refinement, applying a cut-off value of P ≤ 0.05. Genes with Log fold change (FC) values ≥ 1.5 were categorized as up-regulated, those with Log FC values ≤ −1.5 as down-regulated, and genes with Log FC values between −1.5 and 1.5 were considered expressed. Genes meeting the criteria for up-regulation or down-regulation were collectively termed deregulated genes. Detailed statistics for stage-specific and cell line-specific deregulated and expressed genes were compiled in Table S1. Deregulated genes were mapped onto the human protein-protein interaction network (HPPIN) using in-house computational pipelines. This facilitated the construction of three CC stage-specific and two cell line-specific HPPINs for subsequent topological analysis. The deregulated and expressed genes, detailed in Table S1, were subjected to protein-protein interaction (PPI) network analysis, constraining interactions up to the second level interactors. These networks were validated against the random networks as described previously [8, 16]. The detailed statistics of all PPI networks are summarized in Table S2.

### Calculation of IIPs

Important interacting protein (IIP) is broadly classified into four categories, i.e., Hub, CP, GNPP, and LNPP. Genes/proteins satisfying any two of the categories are termed IIP. Hubs are the proteins that have maximum connectivity. Hubs may play a crucial role in the regulation of networks. CP or central proteins are selected based on high betweenness, high clustering coefficient, high closeness, high stress, high ecentricity, and low radiality. GNPP, or global network perturbation protein and LNPP or local network perturbation protein are calculated by performing in-silico perturbation analysis of each node and a node perturbation score is calculated measuring the network centrality parameters of the perturbed and unperturbed network. Perturbation potential of each node was estimated by the global network perturbation score (GNPS) as well as local network perturbation score (LNPS). Careful combination of these network parameters (Hubness, centrality and perturbation potential) led to the identification of crucial nodes for overall integrity of the PPIN [8].

### Cell lines and patient samples

The human cervical cancer cell line HeLa, SiHa, and human embryonic kidney cell line HEK293T were used for the study. All the cell lines were cultured in Dulbecco’s modified Eagle’s medium (DMEM; Gibco) supplemented with 10% fetal bovine serum (FBS-HI; Gibco) and 1% Penicillin-Streptomycin (Pen/Strep; Thermo Scientific) at 37°C in a humidified atmosphere with 5% CO_2_. RNA samples from cervical cancer patients (n=45) were obtained from Tata Medical Center (TMC), Kolkata. Human study (EC/GOVT/15/17) was approved by TMC institutional review board.

### Plasmid constructs and Transfection

The mammalian expression vector pCMV-hAR, harboring the full-length cDNA of human wild type Androgen receptor (AR) (Addgene plasmid # 89078; http://n2t.net/addgene:89078; RRID: Addgene_89078) was procured from Addgene [17]. Corresponding empty vector was generated by removing the AR coding region through BamHI-BglII-mediated digestion. Transient transfections were conducted using both Xfect transfection reagent (Takara) and Lipofectamine 2000 transfection reagent (Invitrogen) following the manufacturer’s protocol. Six hours post-transfection, the culture medium was replaced with media containing 0.1% dimethyl sulfoxide (DMSO) for control cells and with 10nM Dihydrotestosterone (DHT) for pCMV-hAR transfected cells. Cells were maintained for 48 hours post-transfection before initiating each experimental procedure. This standardized incubation period ensured optimal expression and activation of the transfected constructs. For live cell imaging cells were transfected with vinculin-GFP construct 24 hours before conducting the experiments.

### Subcellular fractionation

Cellular disruption and lysis were achieved using an isolation buffer (20 mM Hepes, pH 7.4, 10 mM potassium chloride, 1.5 mM magnesium chloride, 1 mM EDTA, 1 mM EGTA, 1 mM DTT, 0.1 mM PMSF, and 0.25 M sucrose). The lysate was subjected to mechanical disruption by passing through a 25-gauge needle attached to a 1-ml syringe, repeated at least 20 times. This was centrifuged at 600 g to pellet unlysed cell debris. This step was repeated to get the nuclear fractions. The supernatant was further dissolved in immunoprecipitation buffer (50 mM Tris-Hcl, pH 7.5, 150 mM NaCl, 0.1% Triton X-100, 1% IGEPAL, 1 mM PMSF, protease inhibitor cocktail (Sigma Aldrich) followed by TCA and acetone precipitation to get the cytosolic fraction. Subsequently, the obtained fractions underwent analysis through Western blotting using specific antibodies. Antibodies against AR, Lamin B, and GAPDH were employed to probe for the presence of these target proteins in the desired fractions.

### F/G Actin fractionation

Cells were rinsed once in ice-cold PBS before being lysed using Actin stabilization buffer. Cells were kept in Actin stabilization buffer (0.1 M PIPES, pH 6.9, 30% glycerol, 5% DMSO, 1 mM MgSO4, 1 mM EGTA, 1% TX-100, 1 mM ATP, and protease inhibitor) for 10 minutes. Cells were removed by scraping and the whole extract was centrifuged at 4 °C for 75 min at 16,000g. The resulting supernatant, containing G-Actin, was recovered and F-Actin containing pellet was solubilized within Actin depolymerization buffer (0.1 M PIPES, pH 6.9, 1 mM MgSO4, 10 mM CaCl2, and 5 μM cytochalasin D). Following the separation of equal volumes of both suspensions on 12% SDS-PAGE gels, the C4 Actin antibody was used for western blotting. The cellular F/G-Actin ratio was determined by analyzing the optical band density.

### Expression analysis by qRT PCR

RNA extraction from various samples, including local patient samples, HeLa, SiHa, and HEK293 cells, was performed using TRIzol reagent (Thermo Scientific). Subsequently, cDNA synthesis from the extracted total RNA was accomplished using the High-Capacity cDNA Reverse Transcriptase Kit (Applied Biosystems), following the manufacturer’s protocol. RNA expression analysis was performed for local patient samples along with HeLa, SiHa and HEK293cells. Real-time PCR analysis was carried out using PowerUp SYBR Green Master Mix (Applied Biosystems) and gene-specific primers. Relative quantification of each target gene was normalized to the housekeeping gene β-Actin for patient samples and α-tubulin for cell lines and expressed as a fold change compared with untreated control using the comparative cycle threshold (CT) or 2^-ΔΔCT^ method. Gene-specific primers used for the real-time PCR analysis are detailed in Table S5.

### Annexin V-7AAD apoptosis assay

Annexin V-FITC/7AAD staining was employed to assess apoptosis. In brief, cells were seeded at a density of 2×10^5^ cells/well in 6-well plates and transfected with pCMV-hAR or the corresponding empty vector. Ligand addition occurred 6 hours post-transfection. After 48 hours of treatment, cells were trypsinized and resuspended in 100 μl of 1X binding buffer, and stained with 5 μl Annexin V and 2 μl 7-AAD. The cells were gently vortexed, and the mixture was incubated for 15 minutes at room temperature in the dark. Samples were acquired on BD LSRFortessa within 1 h and analyzed with FlowJo software (BD).

### In vitro wound healing assay

Cells were seeded at a density of 5×10^5^ cells/well in 6-well plates. After 24 hours, the cells were transfected with pCMV-hAR and its corresponding empty vector as a control, followed by the addition of the DHT. On the following day, upon reaching confluence, the monolayer was scratched with a sterile 200 μL pipette to create a linear wound in each well. Detached cells were eliminated by washing twice with 500 μL PBS. Subsequently, 1 ml of fresh medium was added and incubated for 48 hours. The progression of scratch closure was monitored and imaged at 0, 24, and 48 hours. Before each image acquisition, the plate was washed with 500 μL pre-warmed PBS to remove debris. Migrated cells were photographed using the EVOS® FL Auto Imaging System (Life Technologies), and the gap area was quantified using ImageJ software.

### Transwell migration and invasion assay

Cells were harvested 48 h post transfection, resuspended in serum-free media and placed in the upper chamber of a Transwell membrane filter (Corning, USA) for migration assays or in the upper chamber of a Transwell membrane filter coated with Matrigel (Corning) for invasion assays. The lower compartment of the chamber was filled with culture medium containing 10% FBS to act as a chemoattractant. Following a 24-hour incubation period, the chambers were extracted, fixed in paraformaldehyde for 30 minutes, and subsequently stained with Giemsa for another 30 minutes. Images were captured, and cell counts were conducted using an Olympus microscope (Tokyo, Japan). Images were captured from a minimum of 9 distinct fields for each condition to calculate the average number of migrated or invaded cells.

### Western Blotting

Whole cell lysates were prepared by dissolving them in 5x loading dye [5% β-Mercaptoethanol, 0.02% Bromophenol blue, 30% Glycerol, Tris-Cl (250 mM, pH 6.8) and 10% SDS] and boiled for 10 min before separating them on SDS PAGE. 10% and 12% Tris-tricine gels were used to separate proteins and transferred to nitrocellulose membranes, which were blocked with blocking buffer (2% BSA or 5% skim milk in TBST). Membranes were incubated overnight with primary antibodies (overnight at 4°C) and the relevant secondary antibodies (2 h at room temperature). Quantification of Western blots was done using GelQuant (Thermo Fisher Scientific). At least 3 independent experiments were performed, and band intensities were normalized to loading control. Details of antibodies and dilutions are provided in Table S6.

### Immunocytochemistry

For immunocytochemistry cells were grown on coverslips and after each experiment cells were fixed with 4% PFA for 10 mins. Permeabilization of cells were carried out using 0.1% Triton X-100 and 10% fetal bovine serum (FBS) in PBS for 1 hour, followed by overnight staining in primary antibodies at 4°Cin a humid chamber. The anti-AR and anti-Vinculin antibody was used at a dilution of 1:100. On the next day, cells were washed in wash buffer containing 10% FBS in PBS and then incubated with secondary antibody solution for 2 hours. Actin filaments were stained with Phalloidin for 30 mins. Coverslips were then washed thrice with wash buffer and observed under confocal microscope.

### Measuring cell movement through time-lapse video microscopy

Cells grown in 35-mm clear-cover glass-bottom confocal dishes (SPL Lifesciences) were transfected with empty vector (EV) and AR followed by DHT treatment (DHT-AR). Post 24 hours of transfection, cells were further transfected with Vinculin-GFP plasmid and kept for another 24 hours. Both the EV and DHT-AR transfected cells were imaged in CO_2_ independent media maintaining conditions of live-cell imaging. Live-cell imaging was performed using ZEISS LSM710/ConfoCor 3 with a 63×1.4 NA oil immersion objective. Movement of an individual cell was imaged for 50 minutes, with an interval of one frame per minute, at room temperature.

### Confocal imaging and image analysis

Confocal imaging of immunocytochemistry samples was conducted using ZEISS LSM 980 and Nikon A1R + Ti-E with N-SIM microscope systems. An Ar-ion laser (for excitation of GFP or Alexa Fluor 488 at 488 nm), a He-Ne laser (for excitation of Alexa Fluor 546 at 543 nm) and a He-Ne laser (for excitation of Alexa Fluor 633 at 633 nm) were used with a 100×1.4 NA oil immersion objective. Z-stacks with a z-interval of 0.15 μm were captured, and initial image analyses, along with 3D projections, were performed using ImageJ. For quantitative analysis, 5 z-stacks from at least 7 images of each replicate were acquired, and this process was repeated across 3 independent experiments. Movement of an individual cell was tracked and analyzed manually in ImageJ.

### Quantitative analysis of focal adhesion size

3D confocal microscopic images of cells with a resolution of 1024 x 1024 pixels (85.33 μm x 85.33 μm) were utilized for quantitative image analysis. In these images, vinculin was stained with secondary antibody (tagged with Alexa Fluor 488) to observe focal adhesions. The green channel images were processed separately using MATLAB’s image processing toolbox to identify the shape dynamics e.g., area and length of vinculin puncta under controlled (EV) and treated (DHT-AR) conditions [18]. Each raw green channel image was initially sharpened using a Gaussian lowpass filter. The sharpened images were then undergone median filtering where each output pixel contains the median value in a 3-by-3 neighborhood around the corresponding pixel in the input image. Lastly, Otsu’s method was used to choose a global threshold from those processed grayscale images for minimizing the intra-class variance of the thresholded black and white pixels. Those thresholds were used to binarize the images to isolate the vinculin puncta. We filter out noisy puncta (< 0.25 μm^2^) using a morphological operation opening. Finally, MATLAB inbuilt function ‘*regionprops*’ was used to calculate area and length (Major Axis) of vinculin puncta [19].

### Statistical analysis

All experiments were conducted with a minimum of triplicates, and statistical analyses were carried out using GraphPad Prism Software version 8 (GraphPad, San Diego, CA, USA). To determine statistically significant differences between control and experimental groups, an unpaired, 2-tailed Student’s t-test was employed. The results are presented with standard error bars indicating mean ± SEM, and the significance of the values is denoted as *P≤0.05, **P≤0.01, and ***P≤0.001.

## Results

### PPIN analysis identified AR as a potential regulator in cervical cancer

The comparative analysis was performed across all cervical cancer stages and cell lines present in the publicly available dataset GSE9750, aiming to identify master regulators potentially implicated in disease development and progression. By combining cervical cancer stage-specific deregulated and expressed genes, disease-specific protein-protein interaction networks (PPINs) were constructed [Table S1, S2]. Subsequent network analyses across all stage and cell line specific networks unveiled 25 Hubs, 1 Central Protein (CP), 4 Global Network Perturbation Proteins (GNPPs), and 4 Important Interacting Proteins (IIPs) consistently present across all stages and cell lines [Table S3]. From this pool, a final set of 11 genes, comprising cell cycle regulators, proliferation markers, proteins involved in protein folding, and steroid hormone receptors, was selected for experimental validation [Table S4]. Among them AR emerged as a critical hub with maximum connectivity displaying downregulation in advanced disease stages. Flowchart of the CC stage specific and cell line specific deregulated network analysis and subsequent validation of the expression of identified genes are outlined in Figure 1A.

**Figure 1:**
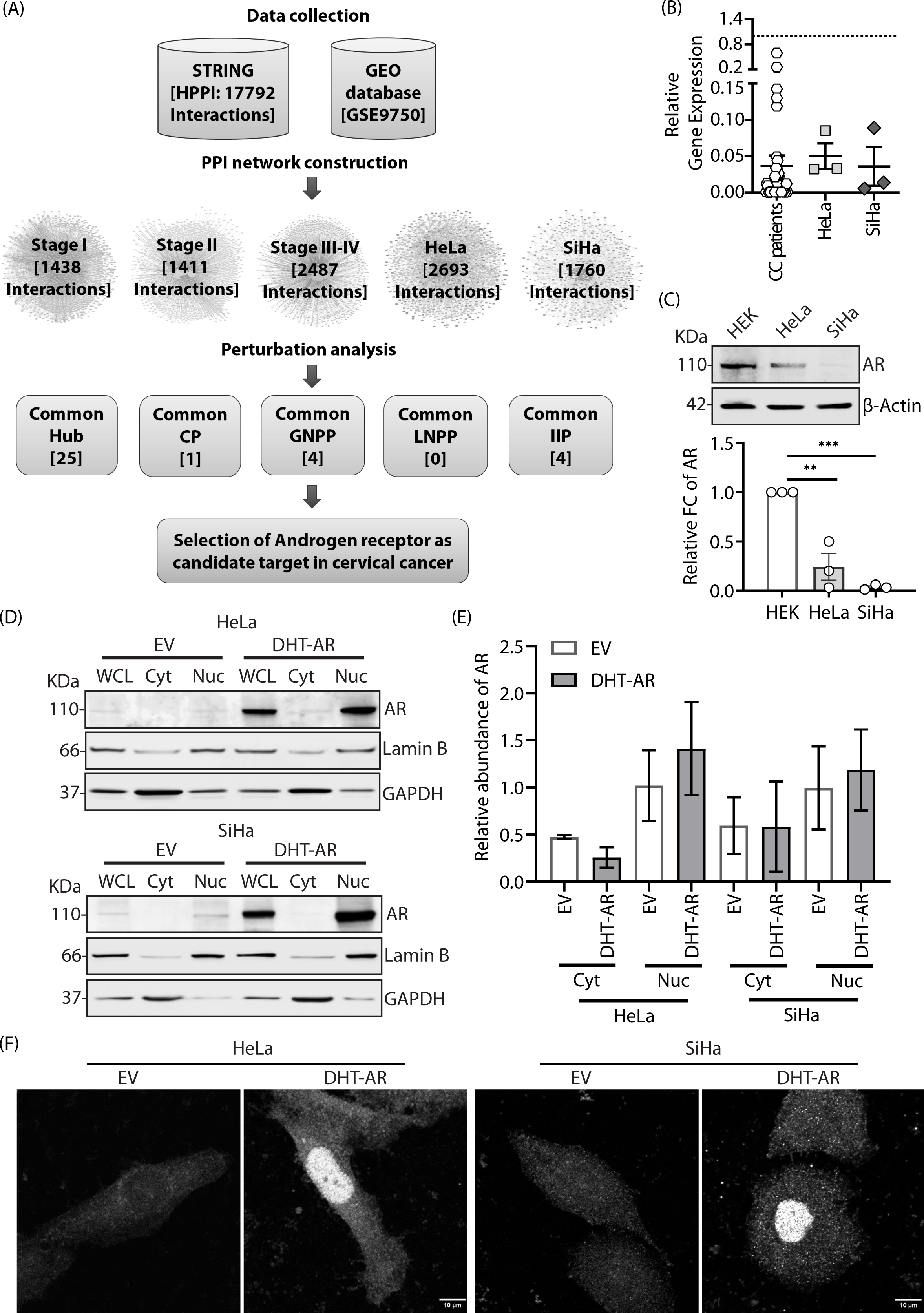
Cervical cancer (CC) specific PPIN analysis and estimation of AR abundance in CC patients and cell lines. (A) Flowchart and steps of cervical cancer specific protein-protein network analysis driven identification of Androgen receptor as critical interacting protein. (B) Relative gene expression of AR in cervical cancer patients, HeLa and SiHa cell lines. mRNA expression of AR was normalized against α-tubulin. (C) Protein expression of AR in cervical cancer cell lines respect to HEK293T cell line. β-Actin was used as a protein loading control. (D) Immunoblotting analysis of cytoplasmic and nuclear distribution of endogenous and overexpressed AR in presence of ligand, DHT for both HeLa and SiHa cell lines. Lamin B and GAPDH were used as markers for nuclear and cytoplasmic fractions, respectively. (E) Quantification of the subcellular localization of AR. Bar graph shows the ratio of cytoplasmic or nuclear fractions with respect to the whole cell lysate of AR. (F) Nuclear localization of AR as detected by immunostaining with AR antibody in both the cell lines, HeLa and SiHa. Scale bar, 10 µm. Data are represented as the mean ± SEM (n = 3). **P < 0.01, ***P < 0.001.

### AR is downregulated in cervical cancer patients and cell lines

Expressions of all 11 genes identified before were analyzed for 45 CC patient samples belonging to the local CC patient cohort and in CC cell lines (HeLa and SiHa) [Figure S1]. AR was found to be significantly downregulated in both patient samples and cell lines, consistent with observations in the transcriptomics dataset [Figure 1B]. Protein expression of AR was also found to be downregulated in the cell lines when compared with the control [Figure 1C]. Taken together, these suggested a potential role for AR in cervical cancer, warranting further investigation into its functional implications and downstream effects in disease progression. Overexpression and ligand (DHT)-induced receptor activation led to enrichment of AR in the nucleus as assayed by subcellular fractionation studies in CC cell lines, when compared with the controls [Figure 1D, 1E]. Imaging studies corroborated these results, where immunocytochemical staining also detected intranuclear localization of AR in DHT treated-AR transfected cells [Figure 1F].

### Exogenous expression of AR followed by ligand activation did not induce apoptosis

To evaluate the impact of AR in cervical cancer, AR was exogenously expressed in the indicated cell lines and 6 hours post-transfection DHT was added for receptor activation; the control cells were treated with equivalent concentration of DMSO. Cells were then stained with Annexin V and 7-aminoactinomycin D (7AAD) to analyse the apoptotic cell population using flow cytometry. FACS data did not show a significant effect of DHT-AR on apoptosis in across samples [Figure 2A, 2B]. This finding was further supported by unaltered protein levels of cleaved and total CASP3 and CASP9 [Figure 2C, 2D].

**Figure 2:**
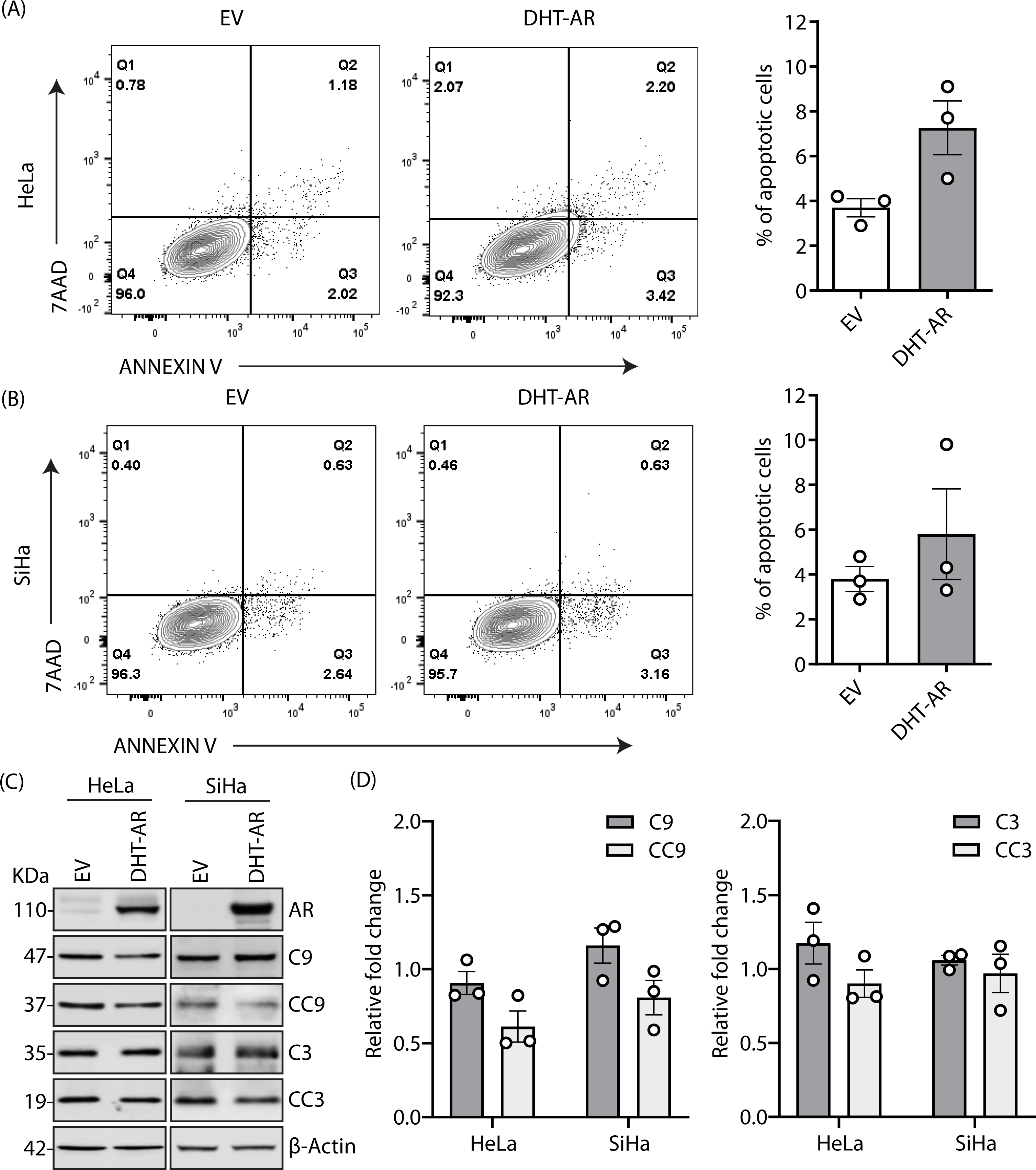
Impact of exogenous expression followed by ligand activation of AR on apoptotic properties CC cell lines. (A) Representative contour plots of Annexin V/7-AAD dual staining to detect the effect of AR overexpression and ligand mediated receptor activation on apoptosis of CC cell lines. (B) The bar graph represents the Annexin V and 7AAD positive cells. (C) Immunoblot of AR, Caspase 9 (C9), cleaved Caspase 9 (CC9), Caspase 3 (C3) and cleaved Caspase 3 (CC3) in HeLa and SiHa cells in empty vector (EV) and AR transfected (DHT-AR) cells. β-Actin was used as a protein loading control. (D) Bar graph represents relative fold change of protein expression where cleaved caspases are normalized to total caspase levels and total caspases are normalized against β-Actin. Data are represented as the mean ± SEM (n = 3).

### Ligand-induced AR activation AR altered cellular migration

Scratch assays were performed to investigate the influence of DHT-AR on two-dimensional movement of CC cells. A decrease of ∼50% was observed in the wound healing capability of DHT-AR cells compared to the controls [Figure 3A]. Additionally, to investigate whether either of the treatment components could elicit a similar effect alone, cells were either transfected with AR or treated only with DHT and subjected to similar scratch assays in HeLa cell line [Figure S2A]. Only AR overexpressed cells (DMSO-AR) were unable to exhibit any significant response on wound healing [Figure S2A]. The ligand (10nM DHT) alone failed to elicit significantly different wound healing capability compared to that achieved in control (DMSO treatment). Further, we used multiple concentrations of the DHT to check if the ligand alone can inhibit the HeLa cell migration. However, we did not observe any significant reduction in migration with higher or lower concentrations of DHT [Figure S2B].

**Figure 3:**
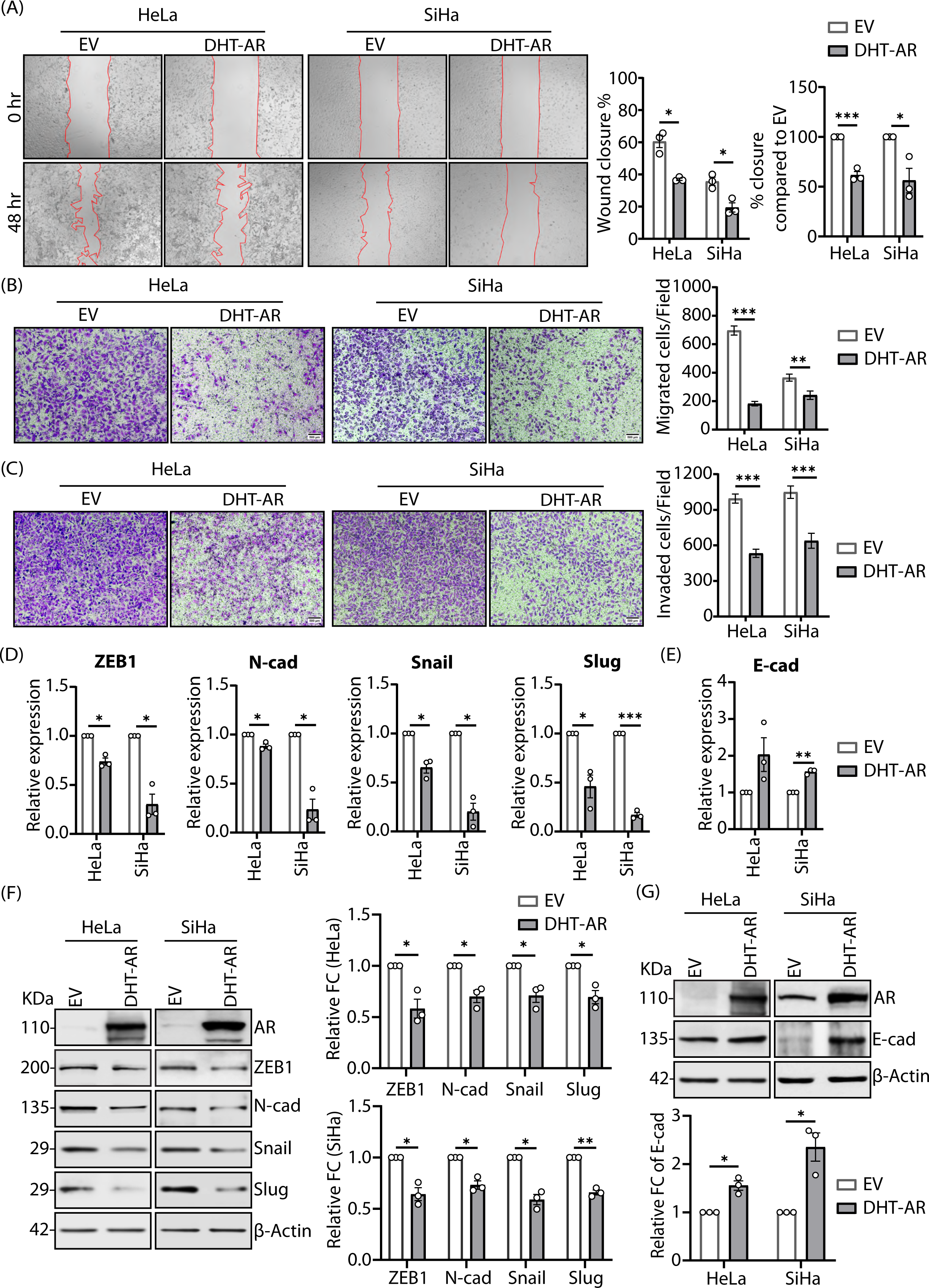
Impact of exogenous expression followed by ligand activation of AR on migration and invasion capability of CC cells. (A) Representative images and quantification of migration using a scratch assay for HeLa and SiHa cells at 0 and 48 hr. (B) Representative images and quantification of migrated cells in transwell migration assay of HeLa and SiHa cells 48 hours post-AR overexpression, in presence of DHT. Field = 29.1 mm^2^. Scale bar, 500 µm. (C) Representative images and quantification of invaded cells in transwell invasion assay of HeLa and SiHa cells 48 hours post-AR overexpression, in presence of DHT. Field = 29.1 mm^2^. Scale bar, 500 µm. (D) Relative mRNA expression level of mesenchymal markers [e.g., ZEB1, N-cadherin (N-cad), Snail, and Slug]. (E) Relative mRNA expression level of epithelial marker E-cadherin (E-cad). (F) Immunoblots and quantification of relative fold change in protein levels of mesenchymal markers in HeLa and SiHa cells 48 hours post-AR overexpression, followed by DHT treatment. (G) Immunoblots and quantification of relative fold change in protein levels of E-cadherin in HeLa and SiHa cells. Data are represented as the mean ± SEM (n = 3). *P < 0.05, **P < 0.01, ***P < 0.001.

Protein levels of established markers for cell migration were verified under similar experimental conditions. Epithelial-Mesenchymal Transition (EMT) orchestrates biochemical changes that enable an epithelial cell to acquire a highly motile mesenchymal phenotype. EMT is crucial for processes like embryonic development and wound healing; however, dysregulation of this process can potentiate metastasis of carcinomas [20, 21]. Hence, the effect of AR on EMT was analysed by Western blotting against two mesenchymal markers, ZEB1 and Snail and an epithelial protein E-cadherin [Figure S2C-S2F]. In presence of DHT, AR-transfected cells showed significantly reduced levels of Snail, when compare with the control; E-cadherin levels were also significantly elevated in these samples. While treatment with DHT or exogenous AR alone did result in an increase in E-cadherin levels, a synergistic effect was seen when both were present. Taken together these findings suggested that in presence of both the factors (DHT and exogenous AR), cell migration could be deregulated. To explore this possibility, cells were subjected to transwell migration assay under the indicated conditions [Figure 3B]. DHT-AR cells exhibited results consistent with those obtained in the scratch assay, indicating a significant reduction in the migratory potential of cells relative to the controls in both the cell lines. To further assess if AR could reduce invasiveness, cells were analysed for their ability to penetrate through matrigel-coated transwell inserts on an invasion assay [Figure 3C]; a significantly low numbers of invading cells per field were detected in presence of DHT and AR overexpressed conditions. Deregulation of EMT markers in turn corroborated these observations, where ZEB1, N-Cadherin, Snail, and Slug were lower at the mRNA as well as protein levels [Figure 3D, 3F]. However, both the mRNA and protein expression of E-Cadherin were elevated in presence of DHT and AR [Figure 3E, 3G]. Our data so far support the hypothesis that AR, in its ligand-activated state could play a role in restricting the migration of cervical cancer cells and prevent invasion.

### PTEN induced GSK3β-mediated degradation of mesenchymal markers

The AR-specific transcriptional regulatory network was extracted from the TRRUST database, with PTEN being a target gene of AR. AR is reported as a transcriptional activator of PTEN in breast carcinoma [22]. We observed 2.5-fold elevated protein levels of PTEN in HeLa cells with AR overexpression and activation [Figure 4A]. However, a similar significant alteration was not detected in SiHa cells. In the canonical PTEN-AKT-GSK3β pathway, PTEN dephosphorylates phosphoinositide-dependent kinase 1 (PDK1), a product of PI3K, thereby negatively regulating the AKT pathway. Reduced phosphorylation of AKT leads to further reduction in the phosphorylation of its substrate, glycogen synthase kinase 3β (GSK3β) [23, 24]. GSK3β, on the other hand, functions as a repressor of mesenchymal markers Snail and β-Catenin by facilitating their proteasomal degradation [25, 26]. In HeLa cells treated with DHT-AR, we observed a significant decrease in phosphorylated PDK1 levels, probably leading to reduced phosphorylation of AKT [Figure 4B]. In cells with DHT-AR, while the protein levels of GSK3β were higher, its phosphorylated form was seen decreased when compared with the control [Figure 4C]. Since reduced protein expression of Snail was observed under similar conditions [Figure 3F], it was obvious to evaluate the status of β-Catenin; its levels were also lower in cells with DHT and exogenous AR, compared with the controls. Snail is a known transcriptional repressor of E-Cadherin [27]. E-cadherin plays a pivotal role in stabilizing β-Catenin at the membrane and sequestering any free β-Catenin that has evaded the cytoplasmic degradation mechanism [28]. As E-cadherin levels were found significantly higher in DHT-AR cells [Figure 3E, 3G]; thus, elevated E-Cadherin expression along with lower Snail and β-Catenin levels suggested a possible involvement of GSK3β mediated degradation of these proteins. It is justified to summarise that DHT-activated AR triggers PTEN activity, this could induce GSK3β-mediated degradation of mesenchymal markers like, Snail and β-Catenin. This would lead to higher abundance of E-Cadherin, which in turn could hinder cell migration [Figure 4E].

**Figure 4:**
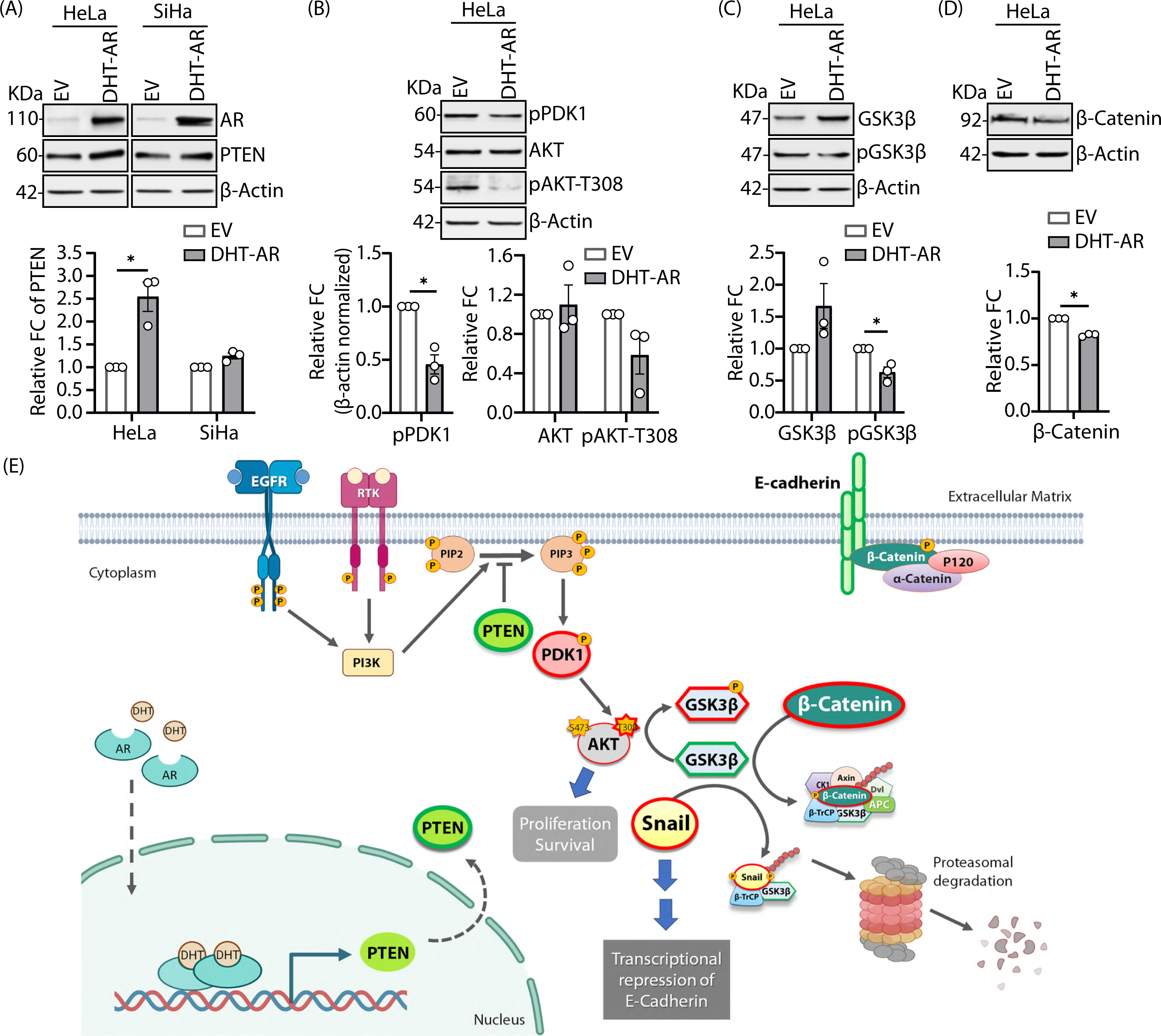
AR-induced regulation of mesenchymal markers. (A) Immunoblots and quantification showing relative fold change in protein levels of PTEN. (B) Immunoblots and quantification of relative fold change of downstream pathway proteins of PTEN involved in the phosphorylation of AKT. Phospho-PDK1 expressions were normalised to β-Actin (C) Immunoblots and quantification showing relative fold change in protein levels of pGSK3β and GSK3β. (D) Immunoblots and quantification showing relative fold change in protein levels of mesenchymal marker β-catenin. Data are represented as the mean ± SEM (n = 3). *P < 0.05. (E) Mechanism of action of AR in restoring GSK3β and downstream functions in HeLa cells via PTEN. Proteins highlighted in bold, green outline are upregulated, while those in bold red outlines are downregulated in AR overexpressed scenario.

### Exogenous expression of AR results in alteration of focal adhesions size

Directional migration of cells requires a simultaneous, synchronized assembly and disassembly of focal adhesions (FAs) at the leading edge of the cell body, alongside their detachment at the rear [29, 30]. To evaluate whether AR overexpression affected f focal adhesions (FA), cells were immunostained against Vinculin and imaged. Significant reduction in the area and length of FAs were observed to be present in cells with DHT-AR [Figure 5A, 5B]. The dimensions of FA size may be used as a predictor of migrational potential of cells [31]. To address this, cells co-transfected with GFP-tagged Vinculin along with control or vector with AR were imaged under live conditions. The AR expressing cells were treated with DHT as before. We observed that in control cells, FAs underwent continuous events of assembly and disassembly; whereas in cells with DHT-AR, they appeared more stationary [Figure 5C, 5D, Movie S1-4]. Quantitative estimation suggested significant reduction of movement in cells with DHT-AR [Figure 5E, 5F]. Vinculin is a key component of focal adhesions and depleted vinculin expression contributes to reduced cell migration and invasion [32, 33]Protein expression of vinculin was found to be significantly low in DHT-AR cells, indicative of the fact of reduced cell migration and invasion [Figure 5G].

**Figure 5:**
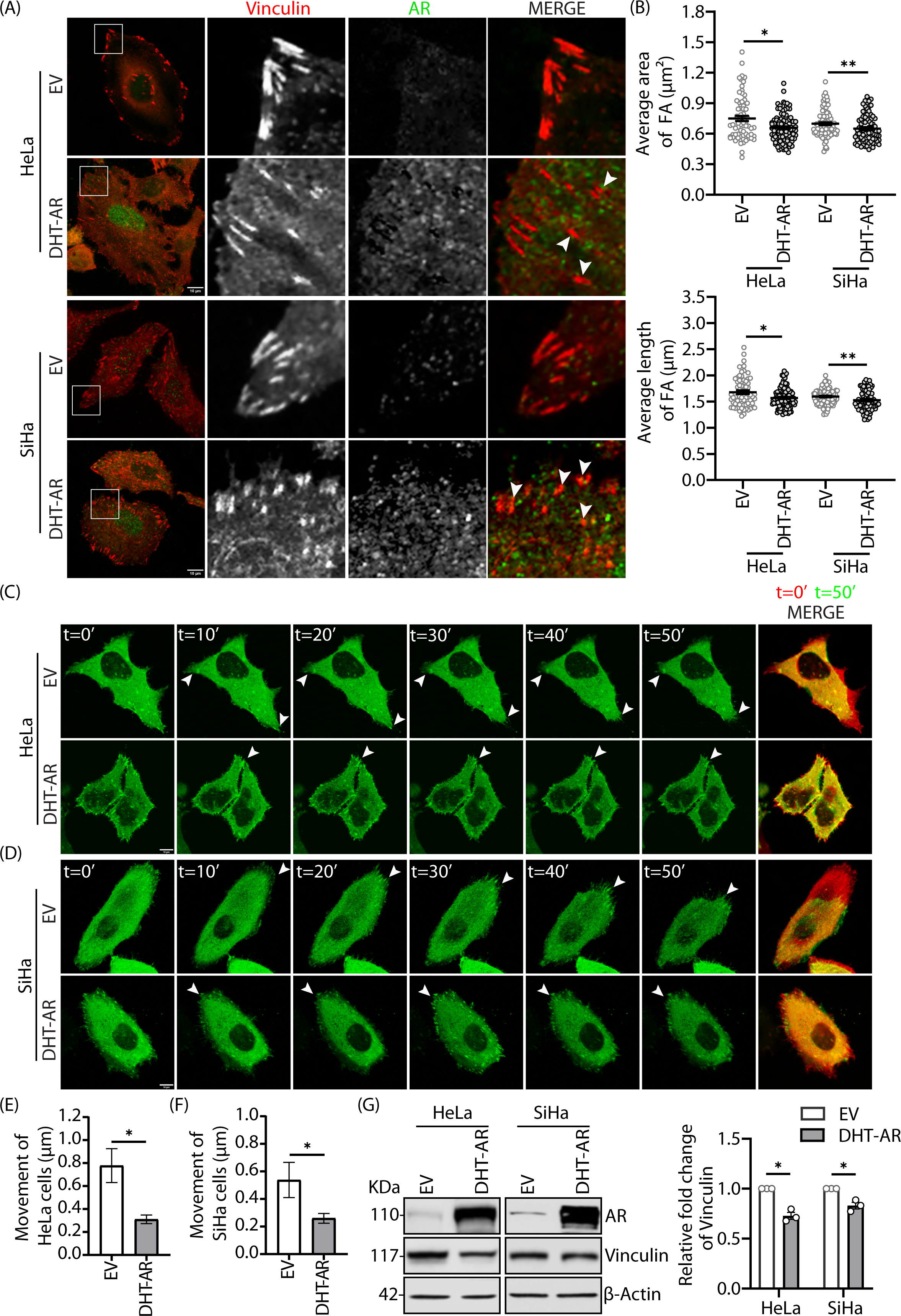
AR overexpression leads to disruption in focal adhesion points in cervical cancer cells. (A) Empty vector (EV) and DHT-AR transfected cells were fixed, co-stained against Vinculin and AR, and imaged for HeLa and SiHa cell lines respectively. Z-stacks (0.15 μm slices) were captured. Images depict a single representative slice. Enlarged views of the areas within the white boxes are shown (insets). White arrowheads mark vinculin puncta representing deformed focal adhesions. Scale bar 10 μm. (B) Scatter plot (up) representative of average area of focal adhesions (FA) and scatter plot (down) representing the average length of focal adhesions. (C-D) Representative time-lapse confocal image stacks of live Vinculin-GFP expressing HeLa and SiHa cells. Confocal image sequence represents position of cell projections at an interval of 10 minutes. White arrowheads mark the cell projections with maximum displacement. Last image in the sequence depicts overlay of cell position at t=0 minute (red) and t=50 minutes (green). The red segments in the overlayed images shows the initial position of the cell and yellow sections represented migrated regions after 50 minutes. Scale bar 10 μm. (E-F) Bar graph represents the quantification of movement of HeLa and SiHa cells. (G) Immunoblots and quantification representing relative fold change of vinculin in HeLa and SiHa cells. Data are represented as the mean ± SEM (n = 3). *P < 0.05, **P < 0.01.

### Exogenous AR disrupts Actin network

Actin filaments are essential for cell motility, and it is widely accepted that disruption of this cytoskeletal network can adversely affect cancer cell migration [34–36]. To address this, cells were immunostained against AR and Actin cytoskeleton was labelled with Alexa Fluor 488-phalloidin. Control cells showed elongated F-Actin fibers uniformly extended across the cell length. In contrast, a punctate distribution throughout the cellular cortex with shorter filament lengths of Actin was seen in DHT-AR cells [Figure 6A]. Since the Actin network was disrupted in the DHT-treated AR expressing cells, we next investigated the status of the RhoA/ROCK1/LIMK1/CFL1 pathway players, the known regulators of Actin network organization and dynamics. DHT-AR cells showed a drastic reduction in protein levels of components of the RhoA/ROCK1 pathway in both the cell lines [Figure 6B, 6C]. Further, the F/G Actin ratio was also significantly decreased in DHT-AR cells [Figure 6D]. Taken together all these results suggest that ligand-activated AR negatively regulates CC cell migration and invasion by plausibly disrupting FAs, in turn regulated by the Actin cytoskeleton [Figure 6E].

**Figure 6:**
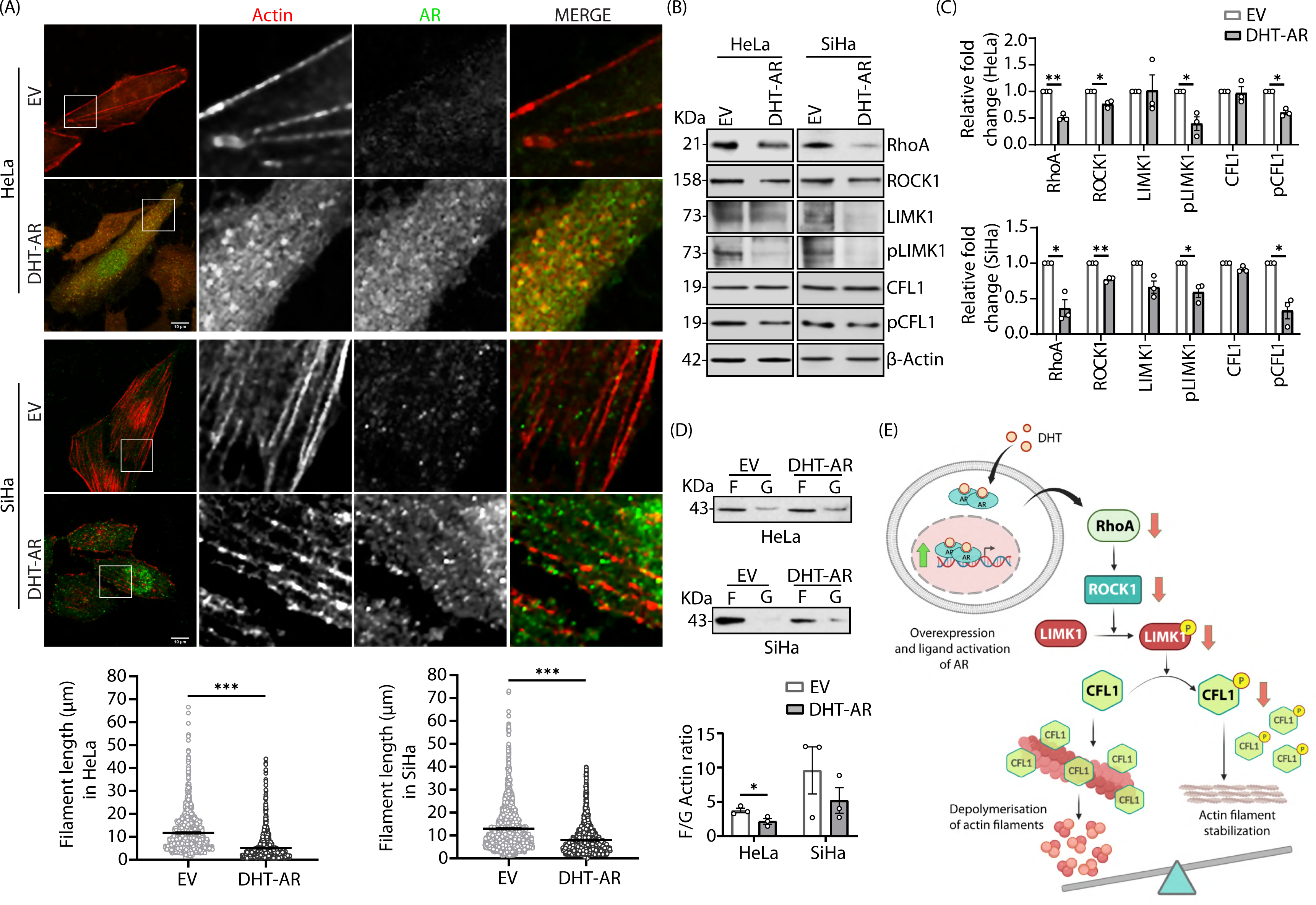
Activated AR destabilizes the Actin network. (A) Empty vector (EV) and DHT-AR cells were fixed, co-stained against AR and Alexa-488 Phalloidin, and imaged for HeLa and SiHa cell lines respectively. Z-stacks (0.15 μm slices) were captured. Images depict a single representative slice. Enlarged views of the areas within the white boxes are shown (insets). Graph shows quantification of Actin length for ∼30 cells from three independent experiments. Scale bar 10 μm (B-C) Immunoblots and quantification representing relative fold change of RhoA/ROCK1/LIMK1/CFL1 pathway proteins responsible for Actin filament formation and maintenance in HeLa and SiHa cells. (D) Immunoblots and quantification representing assessment of Actin polymerization through fractionation to evaluate the ratios of F-Actin (F) / G-Actin (G). Data are represented as the mean ± SEM (n = 3). *P < 0.05, **P < 0.01, ***P < 0.001. (E) Schematic diagram showing AR activation disrupts Actin filament formation by interrupting the RhoA/ROCK1/LIMK1/CFL1 pathway.

## Discussion

Human papillomavirus (HPV) infection is the primary cause of almost all occurrences of cervical cancer, although the disease burden can be substantially alleviated by routine screening and vaccination campaigns. In low and middle-income countries, a huge lacuna remains in the implementation of these simple yet effective measures. A consequence of this imposes the increased risk of deaths due to metastasized cervical cancer in these populations. Hence, in addition to implementing preventative steps, it is imperative to advance our understanding of the pathogenesis to help in our search for better treatment options for advanced disease condition.

Protein-protein interaction networks are essential in providing disease-specific cues that are indispensable for disease development and progression. Our cervical cancer-specific PPIN analysis suggested probable role of AR, which is significantly downregulated. Recently, another study documented a progressive decline in AR expression in cervical cancer patients with the advancement of the disease [11]. Similarly, we detect AR to be highly downregulated in Indian cervical cancer patients, leading us to investigate if altering the state of this hormone receptor could potentially extenuate the severity of the disease. AR being a ligand-activated transcription factor translocates to the nucleus upon ligand binding to initiate transcriptional activation of its target genes. As a master regulator, AR can activate a spectrum of genes that could have either proliferative or antiproliferative functions. Thus, deregulation of AR function may either contribute to disease development or play a protective role. Overexpression followed by ligand activation of AR did not elicit any anti-proliferative response in cervical cancer cells. Rather, overexpression and ligand-mediated activation of AR worked as a negative regulator of migration and invasion in cervical cancer cells. These properties enable cancer cells to detach and relocate from the primary tumor site to distant organs, often facilitated by epithelial-to-mesenchymal transition (EMT). AR overexpression resulted in significant downregulation of classical mesenchymal markers, like, ZEB1, N-cadherin, Snail, and Slug at the mRNA and protein levels. In contrast, E-cadherin, an epithelial marker was significantly upregulated in response to activated AR. This negative regulation of EMT by AR, led us to investigate the probable molecular players participating in this. Analysis of the downstream target genes revealed transcriptional activation of PTEN by AR. Elevated PTEN levels along with reduced phosphorylation of AKT helped restore GSK3β levels. GSK3β activity is essential for maintaining the epithelial architecture, as it facilitates degradation of the downstream proteins responsible for induction of EMT, Snail and β-Catenin. Increased protein levels of GSK3β and reduced expression of Snail and β-Catenin pointed towards AR-mediated repression of EMT markers. Since Snail also works as a transcriptional repressor of E-cadherin, its reduced expression, helped maintain a more epithelium-like phenotype in ligand-activated AR cells. Cell migration is a highly coordinated process that involves physical connections between the extracellular matrix and the Actin cytoskeleton, employing focal adhesions and facilitating signaling between cells and the environment [37]. Ligand-mediated AR activation led to decrease levels of Vinculin protein, smaller and shorter FAs as detected by the presence of Vinculin puncta in cervical cancer cells. It is reported that cell migration speed depends on the adhesion strength between cell and underlying substrates in a biphasic mode [31], suggesting a positive correlation between migration speed and FA size till a threshold. Alteration in FAs got reflected in compromised migration and invasiveness of cells under similar experimental conditions. Further, ligand-mediated AR activated cells exhibited compromised cell movements, unlike the rapidly motile controls.

Apart from FAs, the Actin cytoskeleton is also essential for cell migration and has been reported to influence EMT in metastatic cancer [38]. During EMT, the cortical Actin is rearranged, stress fibers are formed and new Actin-rich membrane-like projections are generated. Ligand activation of AR could effectively destabilize and shorten Actin filaments in cervical cancer cells. RhoA, a key regulator of the cytoskeleton, regulates the rapid polymerization and depolymerization of F-Actin, facilitating the formation of Actin bundles and cellular protrusions [39]. RhoA activation triggers ROCK-dependent remodeling of the Actin cytoskeleton and disruption of E-cadherin based cell-cell adhesions, thereby promoting mesenchymal morphology [38]. ROCK1 is a serine/threonine kinase, which is a downstream effector of the small GTPase RhoA phosphorylates LIM motif-containing protein kinase (LIMK1). LIMK1 phosphorylates the F-Actin-severing protein cofilin (CFL1), inactivating its Actin depolymerization activity and resulting in stabilization of Actin filaments [40–42]. F-Actin generates the motor force for cellular movement and aids in the contraction and retraction of the rear of a cell. This process influences the formation of lamellipodia, playing a crucial role in cancer metastasis [43]. In DHT-AR cells, protein expression of RhoA and its downstream effector ROCK1 were found to be decreased, so were other molecular targets more distal in the pathway. Reduced phosphorylation LIMK1 and lowered expression of phospho-Cofilin 1 (pCFL1) were also detected. It is justified to extrapolate that stabilization of CFL1 protein resulted in subsequent depolymerization of F-Actin. This affected the F/G Actin ratio, phenotypically manifesting as fewer and shortened Actin filaments in cells with reduced motility.

In summary, our study illustrates a holistic approach that integrates PPI networks to identify genes/proteins that play pivotal roles in a specific disease context. AR, being an IIP in cervical cancer specific PPIN, is involved in the regulation of the pathways that can prevent cervical cancer cell migration and invasion by affecting focal adhesion and, Actin cytoskeleton. Our findings underscore the significance of a multidimensional approach combining bioinformatics, network analysis, and experimental validation to identify and characterize critical regulators in cervical cancer pathogenesis.

Currently, the treatment options of cervical cancer mainly involve invasive surgical procedures of removal of uterus or cervix and surrounding structures, followed by radiation or chemotherapy. Hormone replacement therapy (HRT) is being used more often lately due to the severe side effects of chemotherapy. Other than HRT, administration of estrogen or progesterone or both has reduced the risk of invasive cervical cancer [44]. The role of AR in cervical cancer development is one of the least explored areas, though it is more studied in other cancers [45–50]. Women with poly cystic ovarian syndrome (PCOS) and hirsutism have higher than reference values for serum androgen and AR levels [51, 52]. However, PCOS is significantly associated with higher risk of endometrial cancer but does not associate with higher risk of cervical cancers [53]; this supports a probable protectiveness of androgen and AR against cervical cancer development. However, further in depth research is warranted to reveal any such mechanistic connection. Hence, our present study lays out future research direction where DHT or other synthetic androgens along with increased abundance of AR may be explored to reduce and/or delay cervical cancer progression, offering a non-invasive therapeutic option.

## Acknowledgements

SiHa cells were generous gift from Snehasikta Swarnakar (Kolkata, India). We acknowledge Tania Banerjee and Samit Chattopadhyay for processing and providing patient samples. We thank Mrinal K. Ghosh (Kolkata, India) for anti-PTEN antibody and Debashis Mukhopadhyay (Kolkata, India) for providing Anti-Actin antibody [C4]. Vinculin-GFP construct was a kind gift from Dipyaman Ganguly (Kolkata, India). We thank all the members of Oishee Chakrabarti and Dipyaman Ganguly’s laboratories for their constant help and support during the project. SB thanks Priyanka Mallick for experimental support throughout the study.

## Data Availability Statement

The dataset (GSE9750) presented in this study is available from the Gene expression omnibus (GEO) database.

## Author Contributions

SB has performed all the experiments. SD contributed in microscopic image analysis. SM helped in immunocytochemistry experiment. SC, OC, and SB conceived the idea, analysed the data and wrote the manuscript.

