## Supplementary Materials for "Androgen receptor plays critical role in regulating cervical cancer cell migration"

### Supplementary File

**Table S1: Cervical cancer (CC) stage and cell line specific deregulated and expressed genes**

| Patient Stages | Up-regulated Genes | Down-regulated Gene | Deregulated Genes | Expressed Genes |
|---|---|---|---|---|
| **Stage I (n=5)** | 864 | 666 | 1530 | 3622 |
| **Stage II (n=11)** | 593 | 498 | 1091 | 5513 |
| **Stage III & IV (n=11)** | 475 | 831 | 1305 | 6514 |
| **Cell lines** | | | | |
| **HeLa (n=1)** | 1400 | 3847 | 5247 | 1264 |
| **SiHa (n=1)** | 1381 | 2848 | 4229 | 1151 |

**Table S2: Detailed statistics of CC stage and cell line specific PPI networks**

| | No. of Interactions | No. of Nodes | Hub | CP | GNPP | LNPP | IIP |
|---|---|---|---|---|---|---|---|
| **Hela** | 2693 | 1617 | 155 | 197 | 96 | 84 | 42 |
| **SiHa** | 1760 | 1198 | 113 | 150 | 51 | 67 | 30 |
| **Stage I** | 1438 | 938 | 103 | 115 | 54 | 44 | 33 |
| **Stage II** | 1411 | 803 | 79 | 87 | 29 | 24 | 20 |
| **Stage III & IV** | 2487 | 1328 | 135 | 153 | 59 | 50 | 40 |

**Table S3: Common Hub, CP, GNPP, LNPP and IIP in all stages and cell lines**

| | |
|---|---|
| **Hub** | AR, BRCA1, BUB1, BUB1B, CCNA2, CDC45, CDC7, CDK2, CHEK1, EGFR, ESR1, HSPA4, JUN, MCM2, MCM3, MCM4, MCM5, MCM6, MCM7, MDM2, ORC1, PCNA, RAD51, SKP2, SMARCA4 |
| **CP** | LIG1 |
| **GNPP** | BRCA1, EGFR, ESR1, PCNA |
| **LNPP** | - |
| **IIP** | BRCA1, EGFR, ESR1, PCNA |

**Table S4: Genes selected for expression analysis**

| Gene Name | Category | Expression status | | | | |
|---|---|---|---|---|---|---|
| | | **Stage1** | **Stage2** | **Stage3-4** | **HeLa** | **SiHa** |
| **AR** | Hub | Expressed | Expressed | Down regulated | Down regulated | Down regulated |
| **BRCA1** | Hub, GNPP, IIP | Up regulated | Expressed | Up regulated | Up regulated | Up regulated |
| **CDC7** | Hub | Expressed | Expressed | Expressed | Up regulated | Up regulated |
| **DBF4** | Hub | Expressed | - | - | Up regulated | Up regulated |
| **EGFR** | Hub, GNPP, IIP | Expressed | Expressed | Expressed | Expressed | Expressed |
| **ESR1** | Hub, GNPP, IIP | Down regulated | Down regulated | Down regulated | Down regulated | Down regulated |
| **HSPA4** | Hub | Expressed | Expressed | Expressed | Up regulated | Up regulated |
| **HSP90A1** | GNPP | - | Expressed | Expressed | Up regulated | Expressed |
| **MCM2** | Hub | Up regulated | Up regulated | Up regulated | Up regulated | Up regulated |
| **PCNA** | Hub, GNPP, IIP | Expressed | Expressed | Up regulated | Up regulated | Up regulated |
| **SKP2** | Hub | Expressed | Expressed | Expressed | Up regulated | Up regulated |

**Table S5: List of primers used for RT-qPCR detection of mRNAs**

| Gene Name | Forward primer | Reverse primer |
|---|---|---|
| AR | 5’-TCCATCTTGTCGTCTTCGGAA-3’ | 5’-GGGCTGGTTGTTGTCGTGT-3’ |
| ESR1 | GGGAAGTATGGCTATGGAATCTG-3’ | 5’-TGGCTGGACACATATAGTCGTT-3’ |

| | | |
|---|---|---|
| BRCA1 | ACCTTGGAACTGTGAGAACTCT-3’ | 5’-TCTTGATCTCCCACACTGCAATA-3’ |
| HSP90AA1 | ACCTATGGGTCGTGGAAC-3’ | 5’-AGCCTCATCATCGCTTAC-3’ |
| HSPA4 | AGTGATGGATGCAACACAGAT-3’ | 5’-CCAATGTCGTGTCAAATGC-3’ |
| CDC7 | AGTGCCTAACAGTGGCTGG-3’ | 5’-CACGGTGAACAATACCAAACTGA-3’ |
| DBF4 | AGTTGGTAGTGGTGCACA-3’ | 5’-CATCAAATGGACTGCAGG-3’ |
| SKP2 | AGCCCGACAGTGAGAACATC-3’ | 5’-GAAGGGAGTCCCATGAAACA-3’ |
| EGFR | AGGCACGAGTAACAAGCTCAC-3’ | 5’-ATGAGGACATAACCAGCCACC-3’ |
| MCM2 | TGGAGGTGGTACTGGCCA-3’ | 5’-TGACCATGCTGAGCTGGG-3’ |
| PCNA | ACACTAAGGGCCGAAGATAAC-3’ | 5’-ACAGCATCTCCAATATGGCT-3’ |
| ZEB1 | GGCATACACCTACTCAACTACGG-3’ | 5’-TGGGCGGTGTAGAATCAGAGT-3’ |
| N-Cadherin | AGCTCCATTCCGACTTAGACA-3’ | 5’-CAGCCTGAGCACGAAGAGT-3’ |
| Snail | TGCCCTCAAGATGCACATC-3’ | 5’-GGGACAGGAGAAGGGCTTCT-3’ |
| Slug | CGAACTGGACACACATACAGTG-3’ | 5’-CTGAGGATCTCTGGTTGTGGT-3’ |
| α-Tubulin | ACTGGCTCTGGCTTCACC-3’ | 5’-GTCTGAGTGCTCCAGGGT-3’ |
| β-Actin | CACTGGCATCGTGATGGA-3’ | 5’-CCGTGGCCATCTCTTGCT-3’ |

**Table S6: List of antibodies**

| Antibodies | Source | Cat Id. | Dilution |
|---|---|---|---|
| Mouse monoclonal anti-AR | Santa Cruz Biotechnology | SC-7305 | 1:100 (ICC)<br>1:1000 (WB) |
| Rabbit monoclonal anti-Caspase-9 | ABclonal | A18676 | 1:1000 (WB) |
| Rabbit monoclonal anti-Cleaved Caspase-9 | Cell Signaling Technology | 7257P | 1:1000 (WB) |
| Rabbit monoclonal anti-Caspase-3 | Cell Signaling Technology | 9662S | 1:1000 (WB) |
| Rabbit monoclonal anti-Cleaved Caspase-3 | Cell Signaling Technology | 9664P | 1:1000 (WB) |
| Rabbit monoclonal anti-ZEB1 | ABclonal | A21794 | 1:1000 (WB) |
| Rabbit monoclonal anti-N-Cadherin | AbCam | ab18203 | 1:1000 (WB) |
| Rabbit monoclonal anti-E-Cadherin | ABclonal | A20798 | 1:1000 (WB) |
| Rabbit polyclonal anti-Snail | ABclonal | A5243 | 1:1000 (WB) |
| Rabbit polyclonal anti-Slug | ABclonal | A1057 | 1:1000 (WB) |

| | | | |
|---|---|---|---|
| Rabbit monoclonal anti-PTEN | Cell Signaling Technology | 9188 | 1:1000 (WB) |
| Rabbit monoclonal anti-Phospho-PDK1/PDPK1-S241 | ABclonal | AP1357 | 1:1000 (WB) |
| Rabbit monoclonal anti-AKT | Cell Signaling Technology | 4691S | 1:1000 (WB) |
| Rabbit monoclonal anti-Phospho-Akt-T308 | ABclonal | AP1332 | 1:1000 (WB) |
| Rabbit monoclonal anti-GSK3β | ABclonal | A11731 | 1:1000 (WB) |
| Rabbit monoclonal anti-Phospho-GSK3β-S9 | ABclonal | AP1088 | 1:1000 (WB) |
| Rabbit polyclonal anti-beta-catenin | AbCam | ab2365 | 1:2000 (WB) |
| Rabbit monoclonal anti-Vinculin | ABclonal | A2752 | 1:6000 (WB)<br>1:100 (ICC) |
| Rabbit monoclonal anti-RhoA | Cell Signaling Technology | 2117S | 1:1000 (WB) |
| Rabbit polyclonal anti-ROCK1 | AbCam | ab97592 | 1:1000 (WB) |
| Rabbit monoclonal anti-LIM Kinase 1 | ABclonal | A23948 | 1:1000 (WB) |
| Rabbit polyclonal anti-Phospho-LIMK1-T508 | ABclonal | AP0387 | 1:1000 (WB) |
| Rabbit polyclonal anti-CFL1 | ABclonal | A1704 | 1:1000 (WB) |
| Rabbit polyclonal anti-Phospho-CFL1-S3 | ABclonal | AP0178 | 1:1000 (WB) |
| Mouse monoclonal Anti-Actin (Pan) antibody [C4] | ABclonal | ab14128 | 1:2000 (WB) |
| Mouse monoclonal anti-beta-Actin | Santa Cruz Biotechnology | SC-47778 | 1:10000 |
| Mouse monoclonal anti-GAPDH | Santa Cruz Biotechnology | SC-365062 | 1:10000 |
| Rabbit monoclonal anti-Lamin B1 | AbCam | ab229025 | 1:2000 |
| Anti-mouse IgG, HRP-linked Secondary antibody | Cell Signaling Technology | 7076S | 1:6000 |
| Anti-rabbit IgG, HRP-linked Secondary antibody | Cell Signaling Technology | 7074S | 1:6000 |

### Figure S1

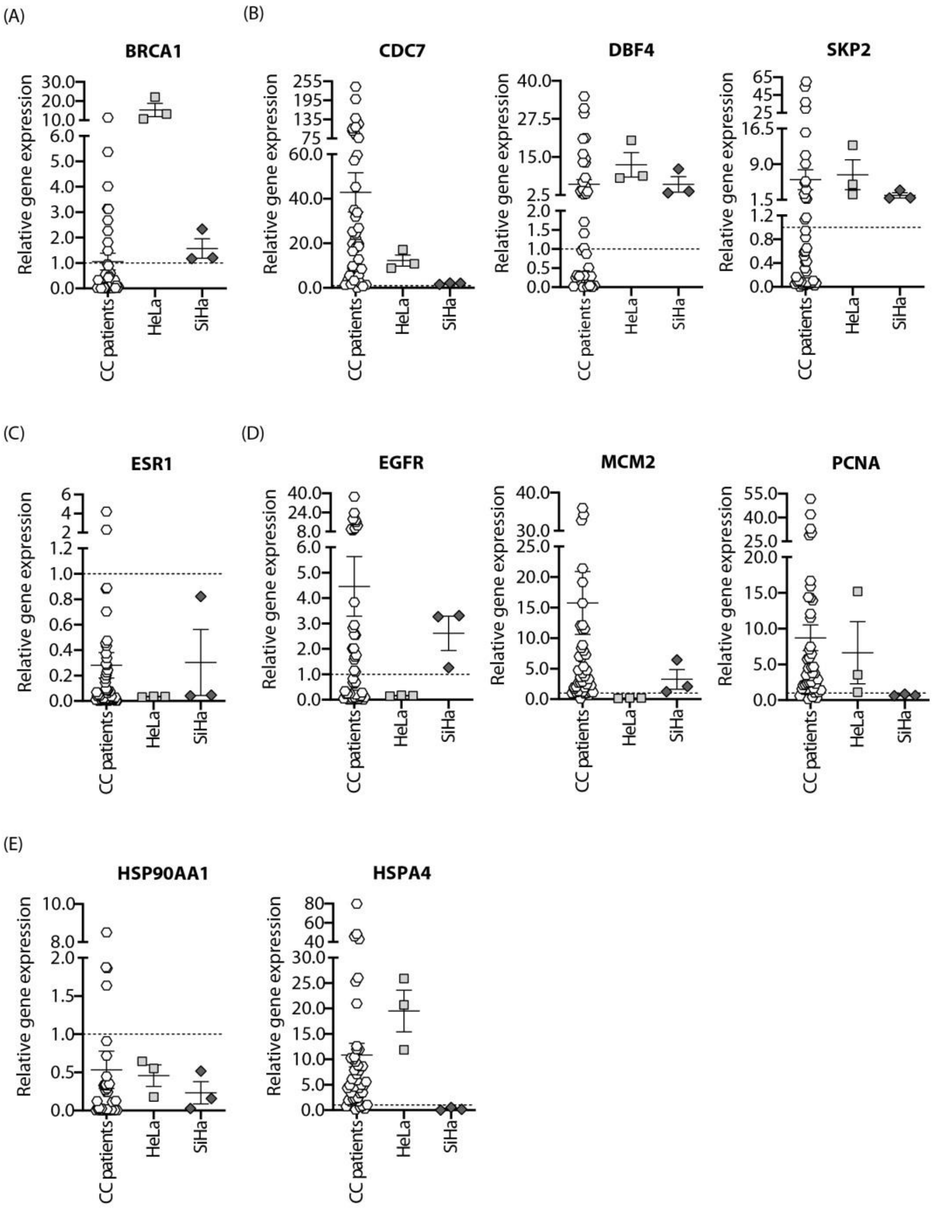


**Figure S1:** Relative mRNA expression of genes selected from network analysis for experimental validation. (A) BRCA1 as tumour suppressor and frequently mutated in cancers, (B) CDC7, DBF4, SKP2 as cell cycle regulators, (C) ESR1 as steroid hormone receptor, (D) EGFR, MCM2, PCNA as proliferation markers, (E) HSP90AA1, HSPA4 as molecular chaperone were selected to be tested in local patient cohort and cell lines of CC.

### Figure S2

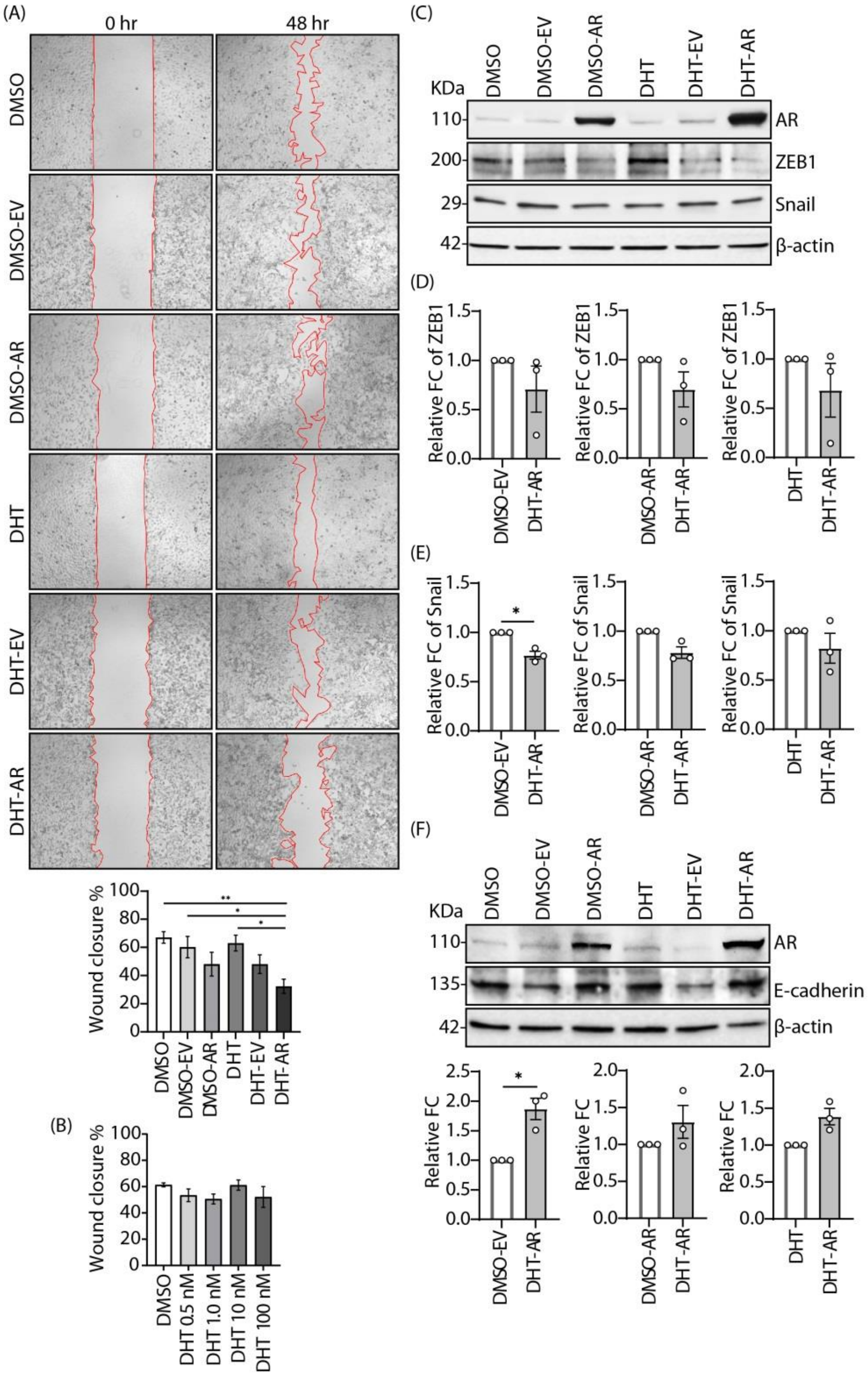


**Figure S2:** Ligand-mediated receptor activation is essential for AR to effectively inhibit the migration of cervical cancer cells.

(A) Representative images and quantification of migration of HeLa cells using a scratch assay at 0 and 48 hr with DMSO (untransfected control), DMSO-EV (empty vector transfected control), DMSO-AR (AR transfection in absence of ligand), DHT (only ligand treated cells), DHT-EV (empty vector transfected and ligand treated control), and DHT-AR (AR transfection in presence of ligand). (B) Quantification of migration of HeLa cells in scratch assay at 48 hr post treatment with different concentrations of DHT (0.5nM, 1nM, 10nM and 100nM). (C-E) Immunoblots and quantification showing relative fold change in protein levels of mesenchymal marker ZEB1 and Snail in DHT-AR cells compared to the ligand (DHT) and only AR transfection (DMSO-AR). (C) Immunoblots and quantification showing relative fold change in protein levels of epithelial marker E-cadherin in the similar conditions. Data are represented as the mean ± SEM (n = 3). *P < 0.05

**Supplementary Movies**

Movie S1: Time-lapse video to track movement of EV transfected HeLa cells.

Movie S2: Time-lapse video to track movement of DHT-AR transfected HeLa cells.

Movie S3: Time-lapse video to track movement of EV transfected SiHa cells.

Movie S4: Time-lapse video to track movement of DHT-AR transfected SiHa cells.
